# Subcellular ToF-SIMS imaging of the snow alga *Sanguina nivaloides* by combining high mass and high lateral resolution acquisitions

**DOI:** 10.1101/2024.07.15.603549

**Authors:** Claire Seydoux, Jade A. Ezzedine, Grégory Si Larbi, Stéphane Ravanel, Eric Maréchal, Jean-Paul Barnes, Pierre-Henri Jouneau

**Affiliations:** Université Grenoble Alpes, CEA, IRIG – Laboratoire Modélisation et Exploration des Matériaux, Grenoble 38000 France; Université Grenoble Alpes, CEA, CNRS, INRAE, IRIG – Laboratoire de Physiologie Cellulaire et Végétale, Grenoble 38000 France; Université Grenoble Alpes, CEA, Leti, Grenoble 38000, France

## Abstract

Time-of-flight secondary ion mass spectrometry (ToF-SIMS) imaging has demonstrated great potential for metabolic imaging, yet achieving sufficiently high lateral and mass resolution to reach the organelle scale remains challenging. We have developed an approach by combining ToF-SIMS imaging acquisitions at high lateral resolution (> 150 nm) and high mass resolution (9,000). The data were then merged and processed using multivariate analysis (MVA), allowing for the precise identification and annotation of 85% of the main contributors to the multivariate analysis components at high lateral resolution. Insights into the electron microscopy sample preparation are provided, especially as we reveal that at least three different osmium-containing complexes can be found depending on the specific chemical environment of organelles. In cells of the snow alga *Sanguina nivaloides*, living in a natural environment limited in nutrients such as phosphorus (P), we were able to map elements and molecules within their subcellular context, allowing for the molecular fingerprinting of organelles at a resolution of 100 nm, as confirmed by correlative electron microscopy. It was thus possible to highlight that *S. nivaloides* likely absorbed selectively some inorganic P forms provided by P-rich dust deposited on the snow surface. *S. nivaloides* cells could maintain phosphorylations in the stroma of the chloroplast, consistently with the preservation of photosynthetic activity. The presented method can thus overcome the current limitations of ToF-SIMS for subcellular imaging and contribute to the understanding of key questions such as P homeostasis and other cell physiological processes.

---

The physiological and developmental functioning of biological organisms relies on their ability to confine specific and optimized biochemical reactions within dedicated compartments at the tissue, cell, organelle and sub-organelle scales. Understanding fundamental processes at the cellular level requires unraveling metabolic processes within the ultrastructural context. Membrane-bound organelles are central in the architecture of eukaryotic cells, each fulfilling several dedicated biochemical functions. While metabolomic profiling of some organelles has been successfully performed for several cell types and organisms (reviewed for plants by Krueger et al.^1^), such approaches are often limited as the process is time-consuming, labor-intensive, and suffers from robustness and reproducibility issues. In addition, for many cell types or species, it is not yet possible. Isolated organelles may not be completely pure, and may be altered during cell fractionation, introducing biases in metabolic analyses.

An alternative to subcellular cell fractionation for spatialized metabolomics is mass spectrometry imaging (MSI), which consists of collecting a mass spectrum from each pixel on a section of an organism, tissue, or cell ^2,3^. The main challenge faced in all MSI workflows lies in the trade-offs between analyte fragmentation, signal strength, acquisition time, lateral and mass resolution ^4^.

Among MSI methods, matrix-assisted laser desorption ionization MSI (MALDI-MSI) is one of the most widely used for biological organisms. Recent developments have improved lateral resolution down to 600 nm^5^, but most current instruments operate at the scale of a few tens of micrometers, which is insufficient for subcellular investigation.

Advances in ion sources for secondary ion mass spectrometry (SIMS) now allow the ionization of large molecules up to 2,000 Da. While the current maximum lateral resolution of ∼ 1 µm is sufficient for single-cell analysis, it remains inadequate for the analysis of most organelles ^6–8^. Harder ionization sources, such as Cs^+^ or O^-^ used on the NanoSIMS instrument, are very effective for subcellular studies, with lateral resolutions of up to 20 nm on biological samples ^9–11^, but limit the analysis to elements or very small molecules due to the high fragmentation induced ^2,12^. Strategies involving isotopic labeling of molecules of interest have been developed ^13^ to mitigate these limitations. An intermediate solution between the ‘harder’ and ‘softer’ sources are small cluster sources such as Bi_3_^+^, available on many current time-of-flight secondary ion mass spectrometry (ToF-SIMS) instruments, which cause less molecular fragmentation but can be focused to submicron lateral resolutions compatible with organelle analysis. These sources have already demonstrated their potential for subcellular imaging ^14–16^.

An additional challenge is that signal intensity in ToF-SIMS becomes lower as the lateral and mass resolutions are higher. These resolutions can therefore only be reached simultaneously at the cost of very long acquisition times.

In this study, we advance ToF-SIMS imaging for subcellular analysis by developing a novel method for data acquisition and analysis. We integrate multiple ToF-SIMS imaging datasets acquired in different modes to obtain high signal intensity, high lateral resolution and high mass resolution from the same sample, pushing forward the capabilities of ToF-SIMS imaging. The datasets are correlated using multivariate analysis (MVA), enabling precise compound identification at high lateral resolution. In addition, we merge them with scanning electron microscopy (SEM) images to identify organelles based on morphological features and thus facilitate metabolic imaging at the submicron scale.

We applied this approach to the study of the snow alga *Sanguina nivaloides*, a unicellular photosynthetic chlorophyte that forms red blooms in melting snow in polar and mountainous regions worldwide. As this alga is not yet cultivable^17^, metabolic studies by subcellular fractionation are not feasible. This limitation makes high lateral and mass resolution mass spectrometry imaging a method of choice to gather spatialized metabolic information on this organism. We demonstrate that multimodal ToF-SIMS/SEM analysis reveals original information related to the sample preparation protocol and provides valuable insights into the metabolism of *S. nivaloides*.

## RESULTS AND DISCUSSION

### Correlative high lateral resolution and high mass resolution analysis

Two blocks of resin-embedded *S. nivaloides* were available from a previous study ^17^. Cell samples collected *in situ* from a snowfield in the French Alps were either *(i)* chemically fixed one hour after sampling using glutaraldehyde and subsequently embedded in resin, or *(ii)* kept at -4°C for a few days before cryofixation and freeze-substitution into the resin. Although cryofixation followed by freeze-substitution best ensures the morphological preservation of microstructures in electron microscopy, the delay in sample handling resulted in significant consumption of lipid droplets and starch compared to the chemically fixed sample^17^. This suggests that the chemistry of the chemically fixed sample remained closer to the native state, so we opted for this block for further analysis. The sample block was sectioned with an ultramicrotome into thin slices (70 nm), which were deposited on a clean silicon substrate.

This sample is complex both from a structural and chemical point of view: the cells contain sub-micrometer scale organelles and various organic and inorganic compounds. Mass spectrometry imaging of these cells therefore requires both high mass resolution to resolve mass interferences between molecules of similar masses, as well as high lateral resolution to resolve small features or organelles. Additionally, it is essential to minimize excessive fragmentation in the high mass range to collect molecular species.

In the NanoTOF II (ULVAC-PHI) ToF-SIMS instrument, the mass resolution depends on the duration of the Bi_3_^+^ pulse, which is in turn bunched to even further reduce its duration. The Bi_3_^+^ beam can also be finely focused to achieve an effective lateral resolution below 100 nm, which requires the use of small apertures. A possible solution to achieve high lateral and mass resolution imaging simultaneously would be to combine a short pulse duration with a finely focused beam. However, this would result in a significant signal decrease, so that an analysis would take several days to reach an acceptable signalto-noise ratio, during which sample lateral drift and stability would eventually become an issue.

To overcome these challenges and achieve both high spatial and mass resolution in a reasonable timeframe, we have therefore developed an alternative approach. It consists of acquiring two independent datasets: one with ultimate lateral resolution and unit mass resolution, and the other with the highest possible mass resolution for the ToF analyzer and a lower lateral resolution. The datasets were acquired sequentially in both polarities using a Bi_3_^+^ beam: high lateral resolution positive and negative (HLR^+^ and HLR^-^) and high mass resolution positive and negative (HMR^+^ and HMR^-^) (Figure 1 A-D), using CGIB sputtering between frames to prevent the accumulation of molecular damage. The HLR acquisitions have a lateral resolution < 150 nm, and the HMR a lateral resolution <500 nm and a mass resolution > 9,000 (Table S1). Molecules were detected in both polarities up to a mass of ∼1,000 (Figure 1 G and 1 H). More than 15 cells, cut in different planes, were imaged in the datasets, giving our findings robustness compared to a single cell analysis. Finally, the positive and negative ion images for each analysis mode were registered and combined two by two (Figure 1 E and F). A representation of the complete workflow presented in this article is available in Figure S1.

**Figure 1.**
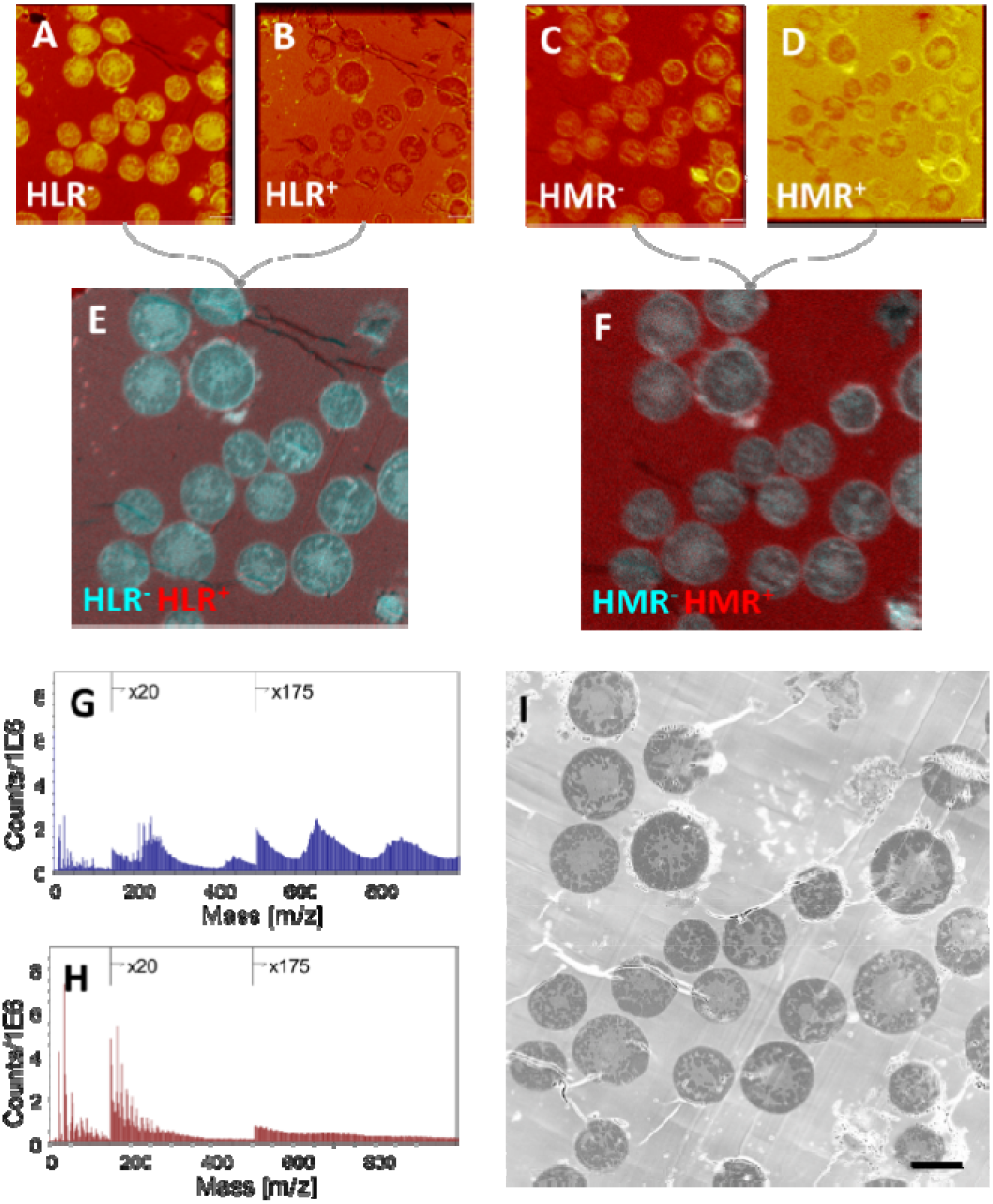
ToF-SIMS datasets at high mass or high lateral resolution and in both polarities and a SEM image were acquired to be further combined. (A-D) Total ion images of (A) HLR^-^ (B) HLR^+^ (C) HMR^-^(D) HMR^+^ were registered and merged into the (E) HLR and (F) HMR datasets. Negative (G) and positive (H) spectra are shown. (I) SEM image over the same area.

The resin sections were cut finely (70 nm) to minimize sample charging, but because dual sputtering consumes material, all four acquisitions could not be performed at the same location. Instead, we chose to use a similar area on two consecutive resin slices, reasoning that the same z-plane would be studied by ToF-SIMS since the width of the slices was much smaller than the lateral resolution. Some differences are, however, visible in the total ion images between the two sets of analyses (Figure 1 A-F), including, for example, folds in the resin at the top right of the HLR images. To assert that the datasets were indeed comparable despite of these small differences, we performed a principal components analysis (PCA) on both datasets separately. The images of the first 5 principal components (Figure S2) are remarkably similar between the HLR and HMR datasets, confirming that most of their variability is due to the same spatial morphological features visible in both datasets. Therefore, they can be compared from a spatial point of view even though they do not physically come from the exact same area.

Furthermore, the datasets were acquired using the same primary ion beam, resulting in similar molecular fragmentation patterns in both modes. The datasets can consequently also be correlated from a spectral point of view. Figure 2 illustrates our strategy for combining the HMR and HLR datasets. The mass resolution in HLR is insufficient to identify any molecule in the ion image (Figures 2 A and 2 B). The corresponding HMR dataset reveals that this peak is composed of Al^+^ (mass 26.9816, mass dev. 0.16) and C_2_H_3_^+^ (mass 27.0239, mass dev. 0.44 mamu, Figure 2 C). As a result, Al^+^ in the HLR image corresponds to the intensity zones around the cells (Figure 2 B and 2 D, red arrows) while C_2_H_3_^+^ shows a diffuse distribution over the observed field of view, with a notably lower intensity in the cell walls and resin folds (Figure 2 B and 2 E, white arrows). The combination of the HMR and HLR datasets thus allows for precise localization and reliable annotation of the molecules.

**Figure 2.**
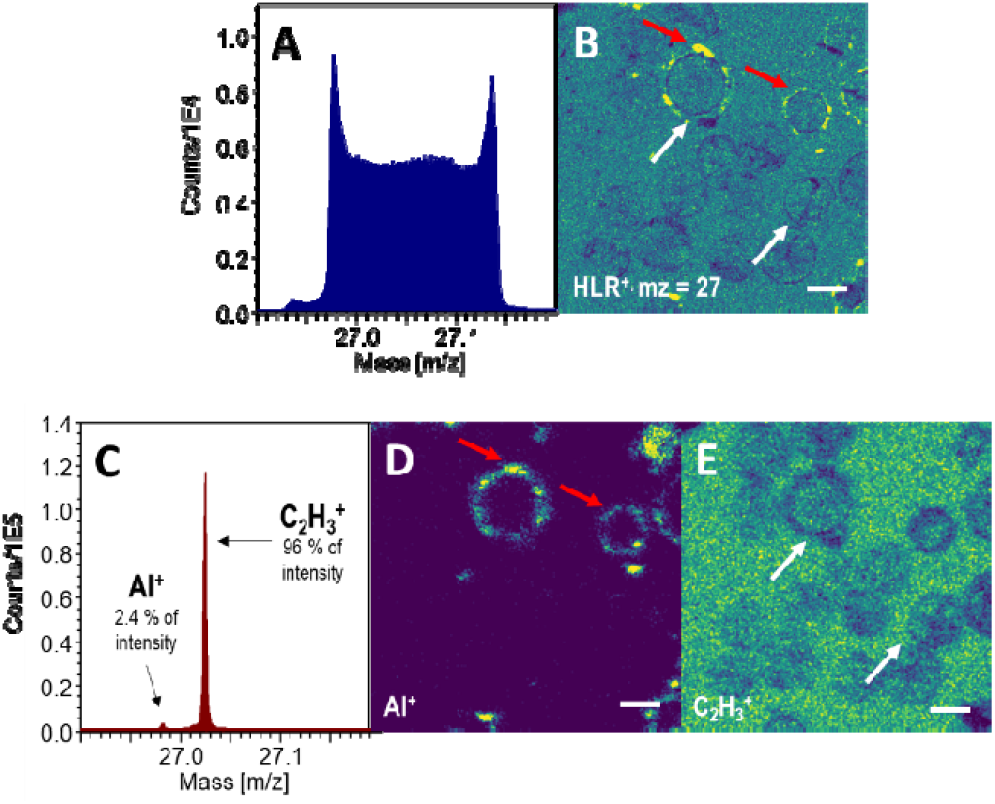
Illustration of the method for HLR and HMR combination. Ion images and spectra for (A, B) HLR^+^ (C, D, E) HMR+ at 27 Da. Scale bars: 10 µm.

Given the complexity of the datasets, the manual identification of all masses using the method described above is not realistic. MVA has proven effective in simplifying complex data, including ToF-SIMS imaging data ^18,19^. PCA is the most popular MVA method, but the spectra of the components contain negative contributions that are difficult to interpret. Another powerful method is non-negative matrix factorization (NMF), also known as multivariate curve resolution. NMF has a positivity constraint and thus decomposes the data into readily interpretable components, providing a basis for segmenting a hyperspectral dataset based on its chemistry.

The HMR and HLR datasets were not merged for NMF analysis. Indeed, due to their different lateral resolutions, their combination would result in components being dominated entirely by either HMR or HLR features and not by a mixture of both. Instead, the HLR and HMR datasets were decomposed separately using NMF (Figure 3, Figures S3 and S4). This approach effectively separates different regions of the image in both datasets– embedding medium, cells, cell walls, chloroplasts, and starch granules.

**Figure 3.**
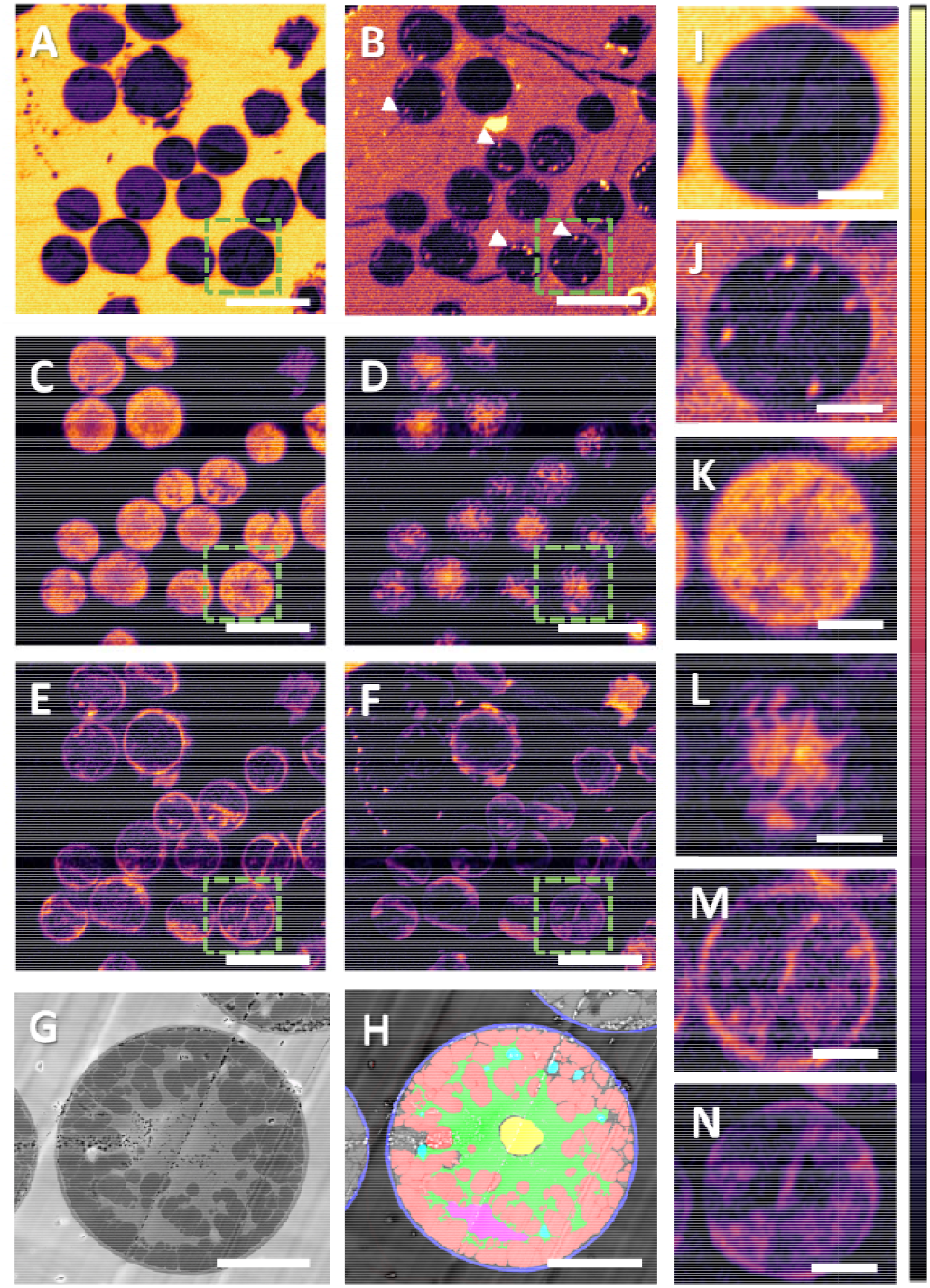
Images of NMF components in the HLR dataset. The images highlight different regions of the sample such as (A) epon resin, (B) resin and starch granules, (C) cell interior and especially lipid droplets, (D) chloroplasts, nuclei and pyrenoids, (E) cell walls and starch granules and (F) cell environment. (G) SEM image and (H) segmentation corresponding to the inset. Red: lipid droplets, blue: cell wall, green: chloroplast, yellow: pyrenoid, cyan: starch granules, magenta: nucleus. (I-N) insets of images (A-F). Scale bars: (A-F): 20 µm, (G-N): 5 µm.

The NMF component images are better defined in HLR mode, but molecules cannot be identified due to the low mass resolution. Therefore, we compared HLR and HMR data, as described previously (Figure 2), to determine which molecules in the NMF spectra contribute to each ion image at a given unit mass. To do this, we assessed which HMR ion image most closely matched the NMF component image. Two levels of identification were sought: identification of the precise mass of the molecule(s) and, when possible, annotation of this molecule. The relative intensity of the identified peak was calculated as a ratio between the identified peak and the total intensity at the unit mass. Using this method, most of the top 20 ions in each of the 6 NMF components were identified (102/120, 85%) and annotated (71/120, 76%) (Table 1, Tables S2 to S6). MVA-assisted correlative analysis thus allows for molecule identification while maintaining the lateral resolution of the HLR dataset. Based on this identification of components, the high-resolution NMR results were further analyzed, revealing the chemical heterogeneity of the sample and providing insights into the fixation and resin embedding procedure.

**Table 1.**
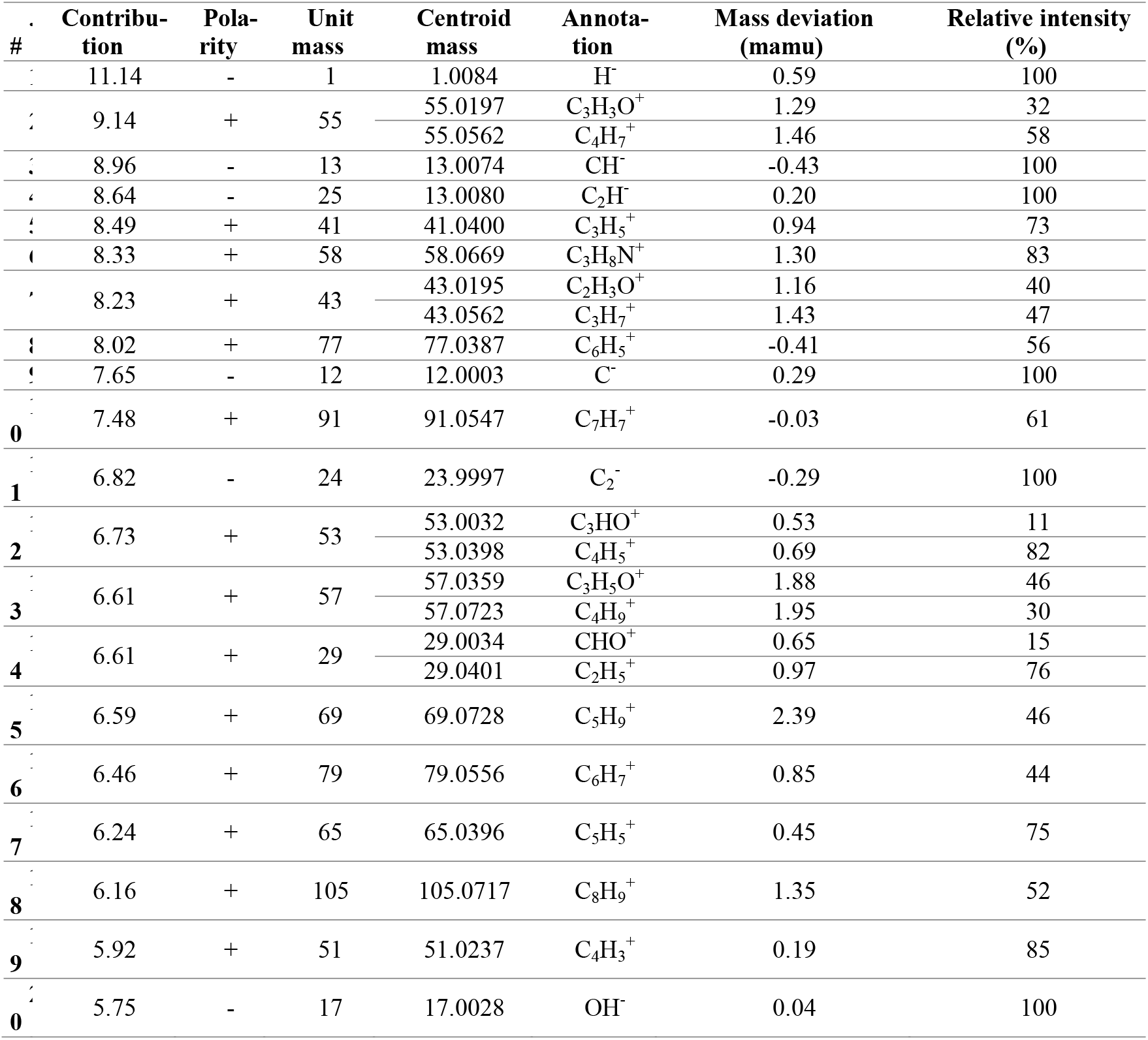
Identification of the top 20 contributors of component A using the correlative HLR/HMR approach

| # | Contribution | Polarity | Unit mass | Centroid mass | Annotation | Mass deviation (mamu) | Relative intensity (%) |
| --- | --- | --- | --- | --- | --- | --- | --- |
| 1 | 11.14 | - | 1 | 1.0084 | H<sup>-</sup> | 0.59 | 100 |
| 2 | 9.14 | + | 55 | 55.0197 | C<sub>3</sub>H<sub>3</sub>O<sup>+</sup> | 1.29 | 32 |
|  |  |  |  | 55.0562 | C<sub>4</sub>H<sub>7</sub><sup>+</sup> | 1.46 | 58 |
| 3 | 8.96 | - | 13 | 13.0074 | CH<sup>-</sup> | -0.43 | 100 |
| 4 | 8.64 | - | 25 | 13.0080 | C<sub>2</sub>H<sup>-</sup> | 0.20 | 100 |
| 5 | 8.49 | + | 41 | 41.0400 | C<sub>3</sub>H<sub>5</sub><sup>+</sup> | 0.94 | 73 |
| 6 | 8.33 | + | 58 | 58.0669 | C<sub>3</sub>H<sub>8</sub>N<sup>+</sup> | 1.30 | 83 |
| 7 | 8.23 | + | 43 | 43.0195 | C<sub>2</sub>H<sub>3</sub>O<sup>+</sup> | 1.16 | 40 |
|  |  |  |  | 43.0562 | C<sub>3</sub>H<sub>7</sub><sup>+</sup> | 1.43 | 47 |
| 8 | 8.02 | + | 77 | 77.0387 | C<sub>6</sub>H<sub>5</sub><sup>+</sup> | -0.41 | 56 |
| 9 | 7.65 | - | 12 | 12.0003 | C<sup>-</sup> | 0.29 | 100 |
| 10 | 7.48 | + | 91 | 91.0547 | C<sub>7</sub>H<sub>7</sub><sup>+</sup> | -0.03 | 61 |
| 11 | 6.82 | - | 24 | 23.9997 | C<sub>2</sub><sup>-</sup> | -0.29 | 100 |
| 12 | 6.73 | + | 53 | 53.0032 | C<sub>3</sub>HO<sup>+</sup> | 0.53 | 11 |
|  |  |  |  | 53.0398 | C<sub>4</sub>H<sub>5</sub><sup>+</sup> | 0.69 | 82 |
| 13 | 6.61 | + | 57 | 57.0359 | C<sub>3</sub>H<sub>5</sub>O<sup>+</sup> | 1.88 | 46 |
|  |  |  |  | 57.0723 | C<sub>4</sub>H<sub>9</sub><sup>+</sup> | 1.95 | 30 |
| 14 | 6.61 | + | 29 | 29.0034 | CHO<sup>+</sup> | 0.65 | 15 |
|  |  |  |  | 29.0401 | C<sub>2</sub>H<sub>5</sub><sup>+</sup> | 0.97 | 76 |
| 15 | 6.59 | + | 69 | 69.0728 | C<sub>5</sub>H<sub>9</sub><sup>+</sup> | 2.39 | 46 |
| 16 | 6.46 | + | 79 | 79.0556 | C<sub>6</sub>H<sub>7</sub><sup>+</sup> | 0.85 | 44 |
| 17 | 6.24 | + | 65 | 65.0396 | C<sub>5</sub>H<sub>5</sub><sup>+</sup> | 0.45 | 75 |
| 18 | 6.16 | + | 105 | 105.0717 | C<sub>8</sub>H<sub>9</sub><sup>+</sup> | 1.35 | 52 |
| 19 | 5.92 | + | 51 | 51.0237 | C<sub>4</sub>H<sub>3</sub><sup>+</sup> | 0.19 | 85 |
| 20 | 5.75 | - | 17 | 17.0028 | OH<sup>-</sup> | 0.04 | 100 |

### Epon resin and osmium components

Components A and B of the HLR analysis (Figure 3 A, B, I, J) and component A’ of the HMR analysis (Figure S4) share a similar location, mainly in the resin. Notably, the spectrum of these components (Figure S3) matches the ToF-SIMS spectrum of bisphenol A diglycidyl ether, also known as Epon, characterized in a previous study ^20^. This is particularly evident with the detection of the peak at mass 211.0758 corresponding to the monomer C_14_H_11_O_2_^-^ (mass deviation 0.80 mamu). The detection of Epon in the sample is expected as the cells were dehydrated and embedded in this resin during the sample preparation.

Component B of the HLR analysis (Figure 3 B and 3 J) shows a similar distribution to component A, except for the presence of small aggregates (∼1 µm in size), corresponding to starch granules as visible in the SEM image (Figure 3 G-H). Some of them are also discernible in the component B’ of the HMR analysis (Figure S4). Our method combining HMR and HLR reveals that this component corresponds to the contribution of carbohydrates C_x_H_y_O_z_^-^ (Table S2), which may indeed originate from the fragmentation of the Epon as well as from the starch located in granules. Identifying these spots as starch granules would have been impossible without the combination of HLR and HMR analyses, especially since the quality of the sample preparation does not allow for visual differentiation between starch granules and mitochondria.

Component C of the HLR analysis is characterized by a periodic signal in the high mass region of the negative spectrum, with a periodicity of ∼200 Da (Figure S3). This periodicity is also visible in the high mass signal of the total spectrum up to mass 1000 (Figure 1G). Furthermore, the main peaks of component C include the isotopic mass distributions of OsO_3_^-^, OsO_2_^-^ or Os_2_O_4_^-^ (Table S4), as determined using the combined HLR and HMR method described previously for peak identification. The periodicity in this spectrum hence corresponds to osmium atoms, incorporated during the sample preparation as a fixative and a contrasting agent to increase backscattered electron signal in SEM. The spatial distribution of osmium oxides (Figure 4A, red) matches that of osmium atoms as evidenced by energy dispersive x-ray spectroscopy (EDX) (Figure 4C) and the electron contrast in SEM (Figure 4B). This confirms that osmium oxides are the primary form of osmium in the sample and as such are responsible for contrast in electron microscopy.

**Figure 4.**
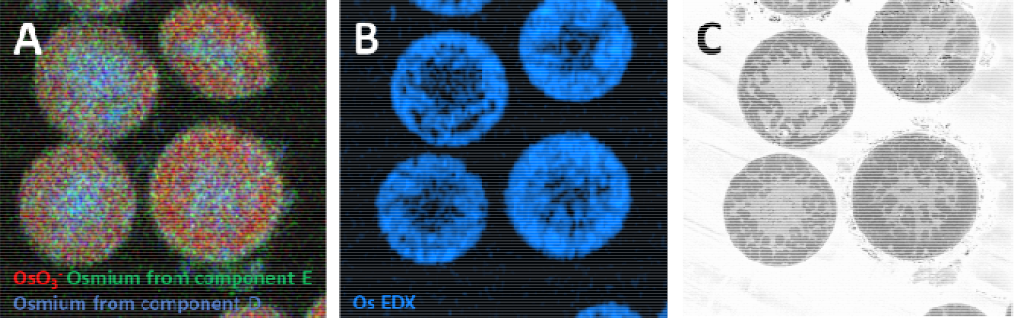
Chemical mapping of osmium. (A) ToF-SIMS: OsO_3_^-^ (red), first four osmium-containing molecules of component E (green), first four osmium-containing molecules of component D (blue); (B) EDX and (C) SEM.

Components D and E also display the same periodicity as component C, indicating the presence of osmium. However, the main osmium-containing peaks surprisingly do not correspond to OsO_3_ or OsO_2_ (Tables S5 and S6), but to other molecules not identified due to the lack of reference spectra in the literature. These molecules differ between components D and E, and their distribution is distinct from osmium oxides (Figure 4A, blue and green). The osmium-containing molecules of component D are mainly located in the chloroplast while those of component E are found in the chloroplast and the cell wall. Additionally, component D is associated with protein fragments in the low mass range. A previous study ^21^ has found that OsO_3_^-^ and OsO_3_(OsO)_n_ ^-^ were differentially localized in adipose tissue. Our analysis suggests that osmium is involved in at least three distinct chemical reactions beyond the esterification of unsaturated bonds in lipids, forming various molecules during sample post-fixation. Osmium tetroxide has been used as a contrast agent and a fixative since the late 19^th^ century for light microscopy and for decades in electron microscopy ^22^, yet the interactions between osmium and biological materials are still not fully understood. Our study provides new insights into the mechanisms by which osmium binds and reacts with biological tissues.

### Molecular profiling of organelles

High mass signals are primarily associated with osmium-containing molecules that are difficult to annotate. However, other significant molecules can be found at masses below 200 Da. Investigating these molecules using the combined HLR and HMR analysis, in correlation with SEM imaging, provided insights on the metabolism of *S. nivaloides*.

Even in HLR mode, the lateral resolution achieved was too coarse to identify the smallest organelles, but correlating spectral images with SEM images overcame this limitation. The backscattered SEM image, with a lateral resolution of 10 nm (Figure 5C), clearly allows segmenting various organelles. Lipid droplets rich in carotenoids and triacylglycerols (orange) fill most of the cytoplasm, and a chloroplast (green), essential for photosynthesis, is visible. At the center of the chloroplast lies a unique pyrenoid (purple), an algal-specific protein-rich compartment where CO_2_ is concentrated (CO_2_-concentrating mechanism, or CCM, involving carbonic anhydrase enzymes) and fixed by the Rubisco protein complex. As the analyzed sample was collected in the field, it is heterogeneous, containing not only algal cells but also bacteria (in red) and dust particles (in yellow). We first focused on nitrogen-containing molecules. Nitrogen (N) is essential for all living organisms, especially as a major component of proteins. Ions such as C_x_H_y_N^+^, CN^-^ or CNO^-^ are considered to be protein fragments ^21,23,24^. These ions are part of component D, which displays protein-related signals (Table S5). Combined ToF-SIMS/SEM analysis shows that they co-localize with bacteria (red arrows), chloroplasts (green arrow), nuclei (pink arrows) and the pyrenoid (purple arrows), with notable intensity in the pyrenoid (Figure 5 A). This high protein density is likely due to the CCM carbonic anhydrase and the CO_2_-fixing enzyme RuBisCo, which are densely packed in the pyrenoid ^25^. In contrast, lipid droplets appear to be poor in proteins. This is consistent with the expected composition of *S. nivaloides* lipid droplets, containing mainly triacylglycerols and carotenoids^17^, and with published data for the green alga *Chlamydomonas reinhardtii*, where similar compartments contain less than 5 % proteins ^26^. Finally, CN^-^ and CNO^-^ ions can also be detected around cells, while C_x_H_y_N^+^ molecules are only found inside cells (Figure 5 A), suggesting that CN^-^ and CNO^-^ do not always originate from proteins. C_x_H_y_N^+^ ions therefore appear more reliable as markers of protein fragments.

**Figure 5.**
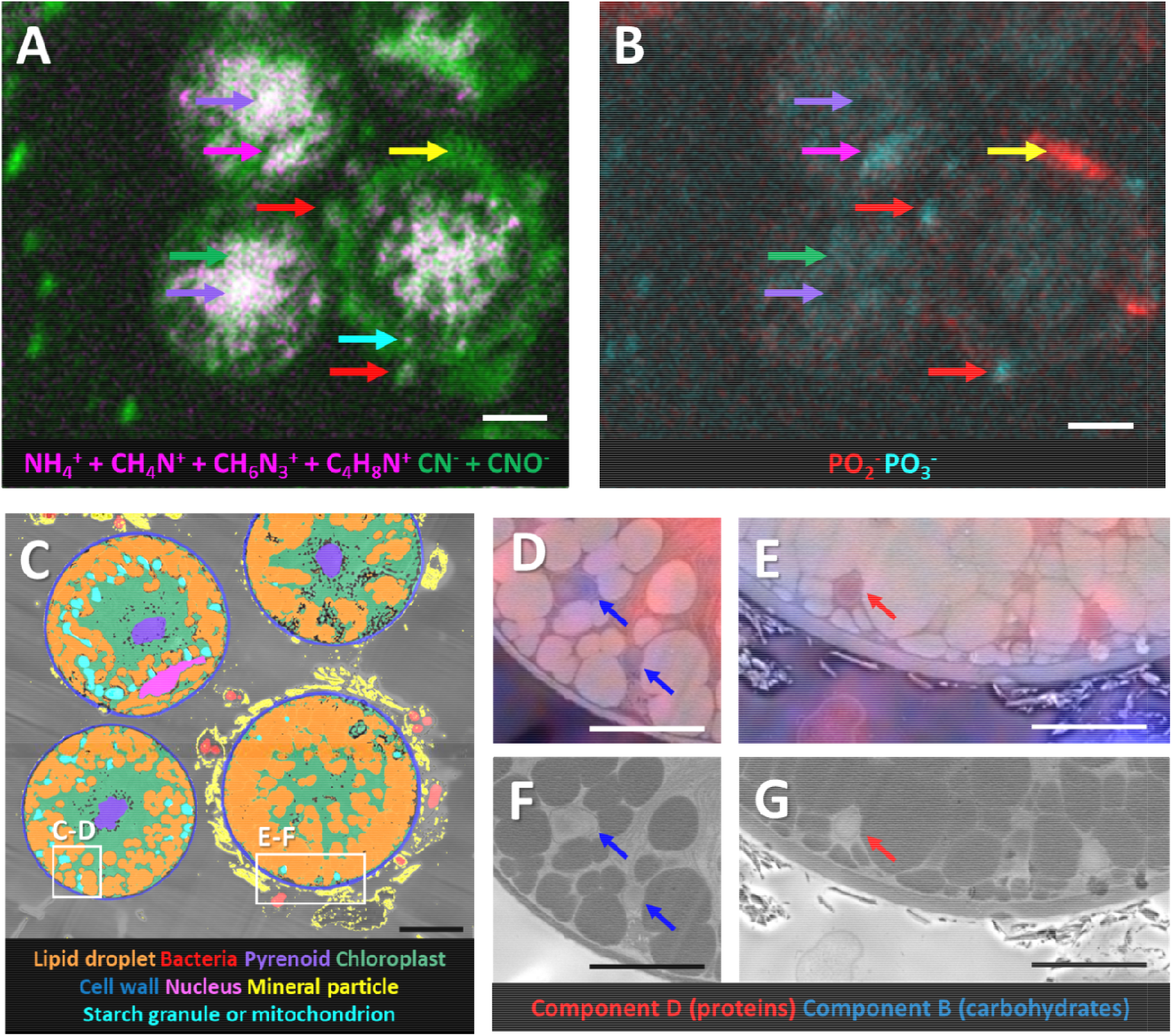
Molecular profiling of organelles in *S. nivaloides*. Ion maps of (A) protein fragments and (B) PO_3_^-^ and PO_2_^-^. (C) Seg SEM image. (D, E) Overlays of NMF components B (blue) and D (red) with subsets of the SEM image in (A) and SEM images s (F) starch granules and (G) mitochondria. Scale bars: (A-C) 5 µm, (D-G) 2 µm.

Phosphorus (P) is another essential nutrient for all biological organisms, serving as a major component of nucleic acids and membrane lipids, as well as contributing to proteins and carbohydrates to varying extents. A molecule often found in ToF-SIMS studies of biological organisms is the phosphocholine ion C_5_H_15_NO_4_P^+^ (*m/*z = 184.1), constituting the polar head of the major phospholipid phosphatidylcholine ^6,14^. A previous study has established that phosphocholine levels were low in *S. nivaloides*^17^, yet we cannot exclude that the absence of the ion is due to a denaturation of lipid species, as the lowering of phosphocholine ion has also been reported in similar sample preparation conditions^21^. We then focused on PO_2_^-^ and PO_3_^-^ ions. PO_3_^-^ is the main peak at its mass (67% of the signal, centroid 79.9580, mass deviation -0.50 mamu). In the HLR analysis C_5_H_3_^-^ (38% of the signal, centroid 62.0230, mass dev -0.42 mamu) strongly interferes with PO_2_^-^ (50 % of the signal, centroid 62.9635, mass dev -0.02 mamu), hence the choice to use the HMR PO_2_^-^ image dataset for this ion.

The P signal was mostly absent from the resin above background noise (Figure 5 C). PO_3_^-^ was detected within the stroma/thylakoids of chloroplasts, which is consistent with the importance of inorganic P as a substrate for photophosphorylation reactions (conversion of adenosine diphosphate, ADP, to adenosine triphosphate, ATP) catalyzed during the photosynthesis process. It could also come from the phospholipids that compose around 10% of the thylakoid membranes. The PO_3_^-^ signal was absent from the pyrenoid (purple arrow). This highlights that, although *S. nivaloides* is starved for this nutrient as previously demonstrated^17^, it has established a subcellular P-saving system, allowing to protect critical activities (photosynthesis and ATP production). PO_3_^-^ was not detected in lipid droplets, consistent with the biochemical composition of this compartment. High PO_2_^-^ and PO_3_^-^ contents were also found in cell nuclei (pink arrows) and in the bacteria surrounding the algae (red arrows). The corresponding profiles show the signature of nucleic acids, which are fragmented into both PO_2_^-^ and PO_3_^-27^ by a Bi_3_^+^ beam. This also suggests that the distribution of PO_2_^-^ and PO_3_^-^ depends on whether the phosphate belongs to a large molecule such as DNA, corresponds to a free inorganic content, or comes from a phosphorylated molecule. In DNA, phosphate groups link ribose residues by two phosphate bonds. Their ionization requires the breaking of two covalent bonds around the phosphate atom, making the formation of a PO_2_^-^ ion more likely than in a molecule where the phosphate groups are more loosely bound.

Interestingly, the PO_2_^-^ zones can be found in the dust particles surrounding the cells (yellow arrow), while PO_3_^-^ is absent from these zones. This suggests that the chemical environment surrounding the phosphate molecules is different, affecting their fragmentation pattern. This may also reflect the selective uptake of environmental phosphorus forms by *S. nivaloides* and bacterial cells, as suggested in a previous study where mineral phosphorus present in the ice was found to be correlated with microorganisms blooms in Greenland glaciers^28^, hinting of a potential mechanism for ice or snow fertilization by mineral dust particles.

Bacteria (Figures 5 A and 5 B, red arrows) are also protein-rich, with higher phosphate-to-protein ratios than algal cell nuclei (pink arrows). This specific content appears to be a distinctive feature of all bacterial cells imaged in the studied datasets, whether encapsulated in mineral particles or not, in the vicinity of algae or free-living, and even across the different species visible in the SEM image. This feature can therefore be interpreted as a chemical signature of the bacteria in the sample.

Finally, while SEM can help to identify the smallest organelles, it does not allow to distinguish mitochondria from starch granules, which have been segmented together (Figure 5C, cyan). Interestingly, components B and D of the NMF HLR analysis, corresponding to carbohydrates and proteins, respectively (Tables S3 and S5), are mutually exclusive in these compartments (Figure 3 B, Figure 5 A, cyan arrow), suggesting that they originate from distinct organelles. In *S. nivaloides* cells, mitochondria have previously been shown to be located specifically in the vicinity of chloroplast protuberances containing starch^17^. Carbohydrates likely originate from starch granules and proteins from mitochondria (Figure 5 D-G). The correlation between ToF-SIMS and SEM therefore allows for a more precise identification of these subcellular structures.

## Conclusion

Biological samples are inherently complex, requiring high mass resolution for metabolic studies by mass spectrometry. Organelle-scale information is essential to understand most biological processes but often missing. The typical size of organelles, bellow or close to 1 µm, challenges the performance of most currently available MSI instruments. Additionally, achieving both high mass resolution and high lateral resolution comes at the expense of signal strength, making their combination in a single analysis virtually impossible. An effective strategy to circumvent these limitations is correlative imaging, where the respective strengths of two techniques can be combined ^23,29–31^. Previous studies have correlated mass spectrometry imaging datasets acquired with two ionization sources, such as MALDI with ToF-SIMS or ToF-SIMS with two different ion beams ^32–34^. These studies often correlate the image of a large molecule at low spatial resolution with an image of one or more putative fragments at higher spatial resolution. Nonetheless, combining these datasets to achieve precise localization of confidently identified molecules is challenging because it requires extensive knowledge of the sample and the fragmentation patterns of the molecules involved using both modalities. To address these limitations, we developed a method for sequentially collecting two separate ToF-SIMS datasets using the same primary ion beam, one with high mass resolution and one with high lateral resolution, and combining them using MVA.

We applied this innovative method to analyze *S. nivaloides* cells prepared by resin embedding. The four datasets (HMR^±^, HLR^±^) were merged into two (HMR and HLR) for MVA. This allowed us to annotate 85% of the most significant molecules in the sub-spectra obtained using NMF, while maintaining a high lateral resolution of 150 nm. Unlike other correlative mass spectrometry imaging approaches using two different modalities, the advantage of our method lies in the correlation of two imaging modes with similar ionization conditions, enabling accurate identification of molecules at high lateral resolution. The development of new machine learning tools ^35–37^ could further improve this method through automatic peak annotation and the creation of ion density maps with higher resolution, but raises a concern about the ability to detect artifacts when they appear.

A limitation of our analysis lies in the sample preparation by resin embedding. Although this method preserves cell morphology and allows for the acquisition of electron microscopy images for ToF-SIMS/SEM correlation, it results in the loss of many large biological molecules such as triacylglycerols and carotenoids hosted in the algal cysts ^17^. Conventional MSI preparations involving cryo-microtomy would significantly damage the sample ultrastructure during freezing, while freeze-drying of the sample is also known to cause uncontrolled delocalization of molecules ^38^, which is incompatible with the desired lateral resolution. An alternative approach to maintain both chemical and morphological integrity would be to perform full cryogenic analyses^39,40^, but these come with further implementation difficulties.

Our analysis provides valuable insights on the preparation protocol, especially regarding the use of osmium. Osmium tetroxide is widely used in electron microscopy as a fixative and staining agent, but its interactions with biological tissues remains debated. Our analysis reveals the presence of three different forms of osmium-containing molecules, each with a unique distribution pattern across cellular compartments. These findings could provide a better understanding of the mechanisms of osmium tetroxide interaction with biological material.

Combining high lateral and high mass resolution ToF-SIMS, correlated with SEM, allows the molecular profiling of organelles or sub-organellar territories such as chloroplasts (including the stroma, the pyrenoid and starch), nuclei or mitochondria. Notably, it highlights the ratio between proteins, PO_2_^-^ and PO_3_^-^ as a chemical signature of specific organelles. Here we show that extracellular P in the snowpack may rely, at least in part, on P-rich dust deposited by winds, and that the uptake of inorganic P by *S. nivaloides* likely mobilizes selective transporters. We also show that the phosphorus-saving mechanism shown previously by the low phosphatidylcholine/betaine lipid ratio in cysts^17^ may not be part of a general degradation of the phosphate-dependent metabolism, in particular in the chloroplast. Rather, our finding suggests that this phosphorus-saving mechanism at the membrane lipid level may well be constitutive to the cyst stage, allowing the cell to keep the capacity to preserve its ATP-dependent energetic metabolism.

The local chemistry of compartments as small as starch granules and mitochondria (< 500 nm) provides sufficient chemical information to identify them, even when their identi-fication through morphology using electron microscopy images fails. All of these findings provide valuable information about the content of cells and may even, in the future, allow for the identification of subcellular structures through their specific chemical signature, without the assistance of electron microscopy. Overall our method demonstrates the capabilities of ToF-SIMS imaging for metabolic profiling of organelles and pushes forwards the capabilities of this method.

## MATERIALS AND METHODS

### Sample preparation

*Sanguina nivaloides* were collected *in situ* from a red snowfield in Vallon Roche Noire (45°02’55.4’’N 6°23’40.8’’E at 2318 m above sea level on June 18, 2021 at 11 in France), chemically fixed and embedded in epon resin as described previously.^17^ The resulting resin blocks were then sectioned into 70 nm thick sections using an RMC ultra-microtome. The sections were collected on clean silicon wafers.

### ToF-SIMS imaging

The wafers were analyzed using a NanoTOF II (ULVAC-PHI) using a 30 keV Bi_3_^+^ beam. Each shot was followed by sample charge compensation using an electron flood gun in both polarities and a 1.5 keV argon gun in the positive polarity. Every second frame from the Bi_3_^+^ gun was followed by a 1 second soft sputtering phase from a 2.5 keV Ar^+^_2500_ clusters from a GCIB gun in non-interlaced mode, followed by a charge compensation phase using both the argon and electron guns. This instrument can operate in two modes: the so-called ‘unbunched’ mode, which provides the highest lateral resolution (HLR) and unit mass resolution, and the ‘bunched’ mode, which allows for high mass resolution (HMR) by shortening the pulse duration at the expense of a weaker signal and lower lateral resolution.

HLR and HMR analyses were performed on the same area on two consecutive resin slices. Because the slice is thinner than the lateral resolution obtained, the same z-plane is studied on consecutive slices. Data were corrected for lateral image drift and mass shift in TOF-DR (TOF-DR 3.2.0.5, ULVAC-PHI). The lateral resolution was also measured in TOF-DR on the total ion image. Relevant peaks were selected manually. Images were exported as .*tif* files for further analysis. All relevant information concerning analysis conditions is presented in Table S1. The complete workflow is shown in Figure S1.

### Scanning electron microscopy

A thin carbon layer (5 nm) was deposited on the wafers after ToF-SIMS analysis to prevent charging during SEM imaging. Images were acquired on the same area on a Gemini SEM 460 microscope (Zeiss) equipped with an aBSD detector (Zeiss), operating at 3 kV and 1 nA. Six images with a pixel size of 9.89 nm were acquired and stitched together using the Image J Pairwise stitching plugin ^41^. The SEM images were then segmented using Ilastik ^42^. The segmentation was afterwards manually curated.

### Image processing

Ion images were convolved using the Kernel 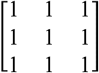 as a means to increase the signal-to-noise ratio without compromising lateral resolution too much. Each image was then normalized to the convolved total ion count image. The SEM image and the total ion images in of the four ToF-SIMS datasets were considered for alignment. Transformation matrices were calculated in Python using the SIFT algorithm implemented in the OpenCV package. Alignments were performed using the StackReg package.

### Multivariate analysis

ToF-SIMS datasets were merged by analysis mode (HLR and HMR) and cropped in the x and y dimensions to the maximum overlap area. Data were then scaled to unit variance and zero mean for principal component analysis (PCA) or to the maximum absolute value for non-negative matrix factorization (NMF). PCA and NMF were performed in Python using the Scikit-learn package ^43^.

### Energy dispersive X-ray spectroscopy (EDX)

EDX analysis was performed using a Merlin microscope (Zeiss), equipped with two Quantax XFlash detectors (Brucker). A spectral image with a resolution of 679 × 658 pixels and a pixel size of 200 nm was acquired. Acquisition was performed at an accelerating voltage of 4 kV, a current of 2 nA, over an acquisition time of 14 h. The various elements detected on the spectra (C-K, O-K, Na-K, Al-K, Si-K, P-K, and Os-M) were then deconvoluted for each pixel of the image using a 2×2 tiling.

## ASSOCIATED CONTENT

### Supporting Information

The Supporting Information is available free of charge on the ACS Publications website.

Supplementary File: Supplementary Figures 1 to 5, Supplementary tables 1 to 6, PDF.

## AUTHOR INFORMATION

### Author Contributions

The manuscript was written through contributions from all authors. All authors gave their approval for the final version of the manuscript.

## ACKNOWLEDGMENT

C. S. and J.-P. B. were supported by the 3D-LIPID project funded by 3BCar and the CEA-Leti Carnot Institute. The authors were supported by the Agence Nationale de la Recherche (GRAL Labex ANR-10-LABEX-04, EUR CBS ANR-17-EURE-0003, Alpalga ANR-20-CE02-0020, PEPR Algadvance A-22-PEBB-0002) and IDEX Université Grenoble-Alpes (Glyco@Alps Cross-Disciplinary Program; Grant ANR-15-IDEX-02). This work was carried out on the Platform for Nanocharacterisation (PFNC), supported by the “Recherche Technologique de Base” and “France 2030 - ANR-22-PEEL-0014” programs of the Agence Nationale de la Recherche.

---

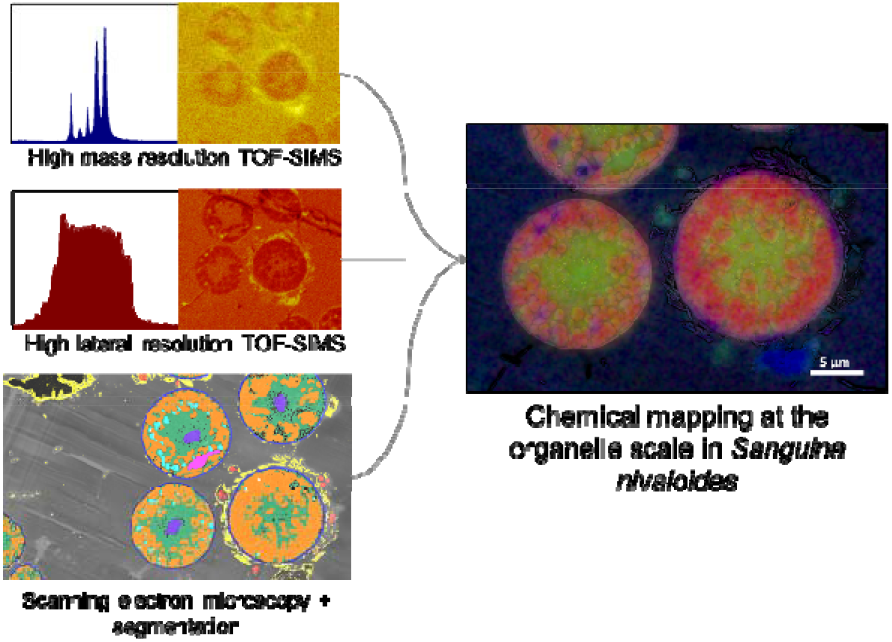

## Notes

### Competing Interest Statement

The authors have declared no competing interest.

### Summary of Updates

Typos in Title, Discussion and Results have been revised.

