## Supplementary data for "Subcellular ToF-SIMS imaging of the snow alga *Sanguina nivaloides* by combining high mass and high lateral resolution acquisitions"

**Table of contents**

**Figure S1.** Workflow used in this study.

**Figure S2.** Spatial location of the 6 first PCA components for the HMR and HLR datasets

**Figure S3.** Images and spectra of NMF HLR

**Figure S4.** Images and spectra of NMF HMR

**Table S1.** Summary of analysis conditions for the 4 ToF-SIMS datasets presented in this study.

**Table S2.** Identification of the top 20 contributors of component B using the correlative HLR/HMR approach

**Table S3.** Identification of the top 20 contributors of component C using the correlative HLR/HMR approach

**Table S4.** Identification of the top 20 contributors of component D using the correlative HLR/HMR approach

**Table S5.** Identification of the top 20 contributors of component E using the correlative HLR/HMR approach

**Table S6.** Identification of the top 20 contributors of component F using the correlative HLR/HMR approach

**Figure S1**

Workflow used in this study.

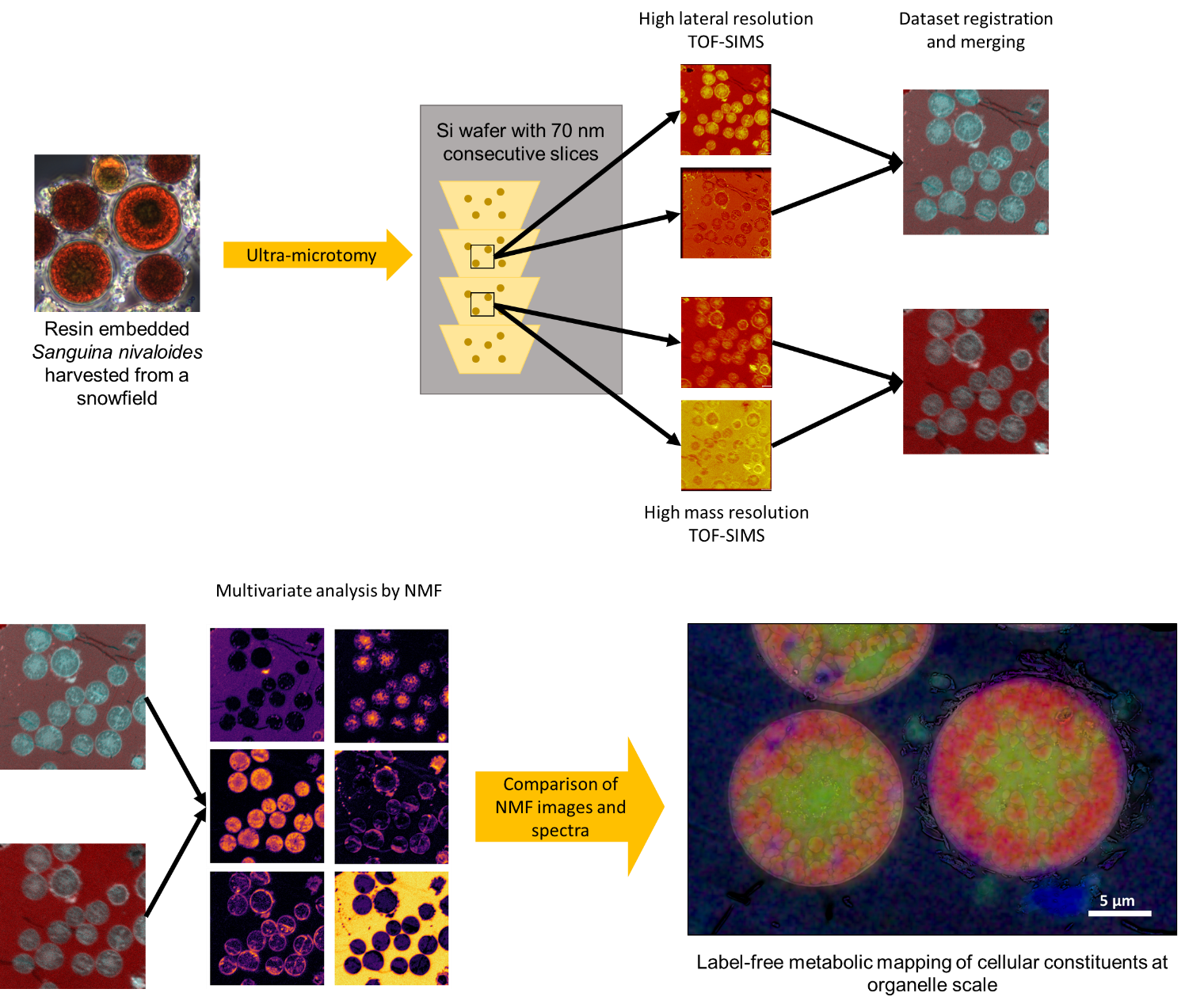

**Figure S2**

Spatial location of the 6 first PCA components for the HMR and HLR datasets.

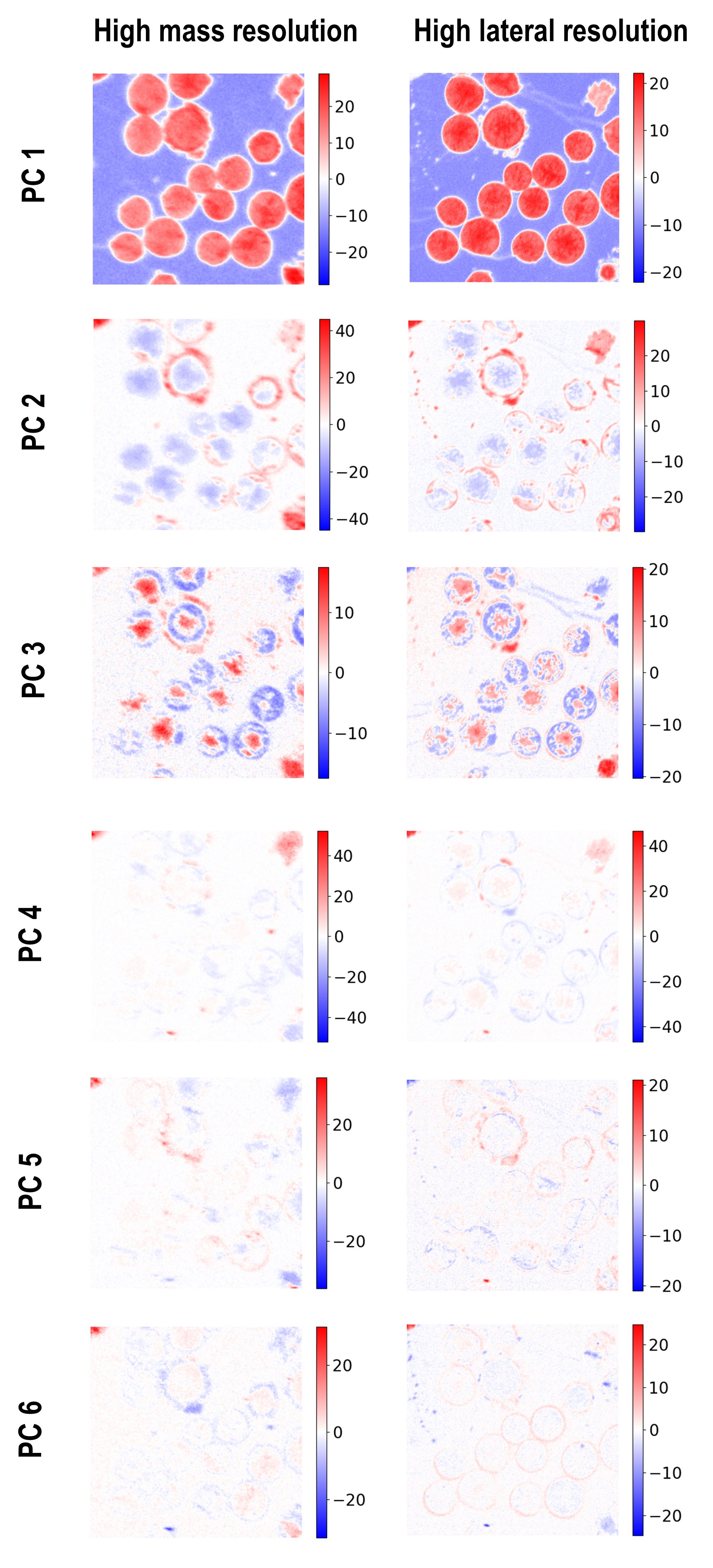

**Figure S3**

Images and spectra of NMF HLR.

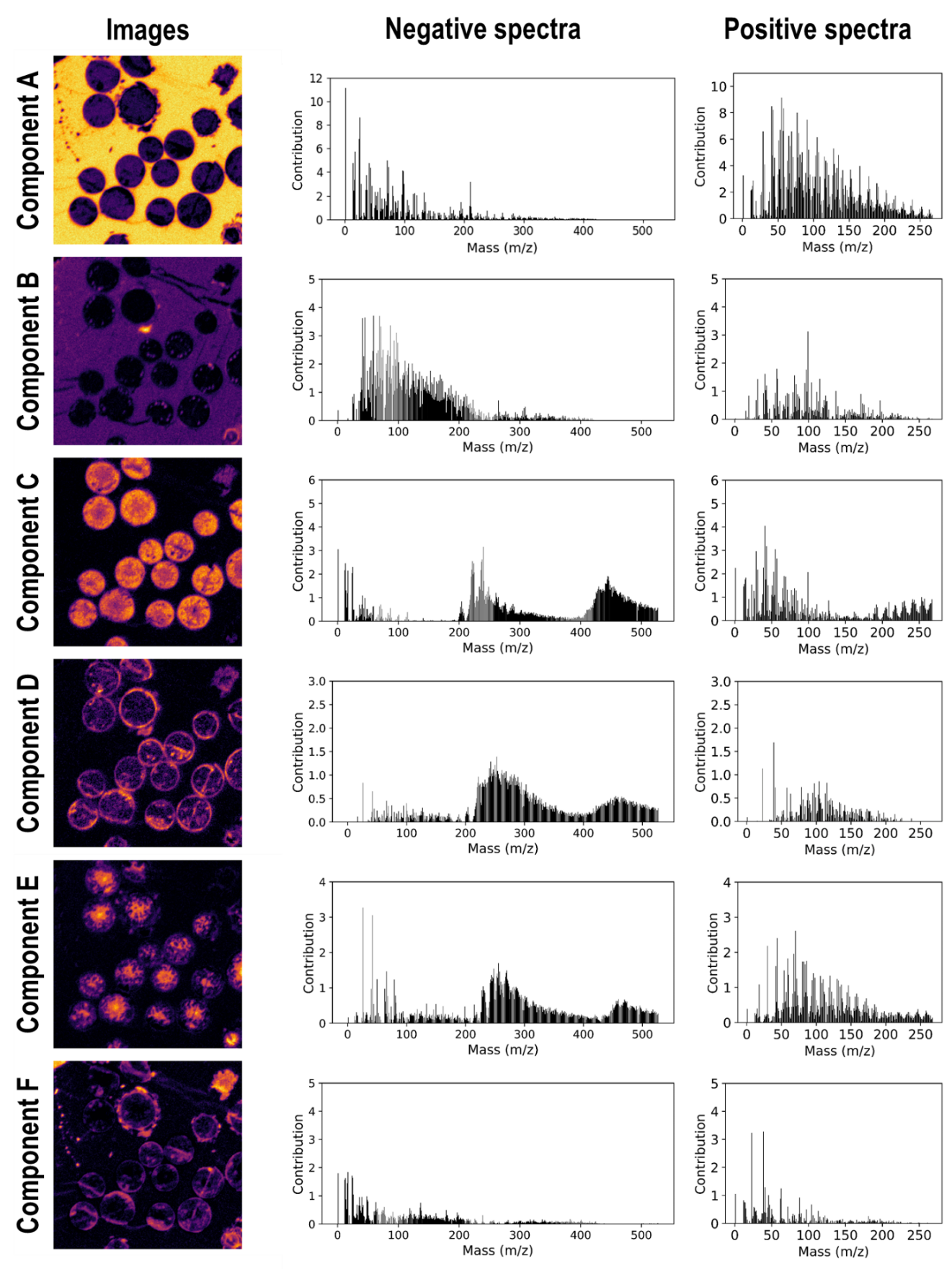

**Figure S4**

Images and spectra of NMF HMR.

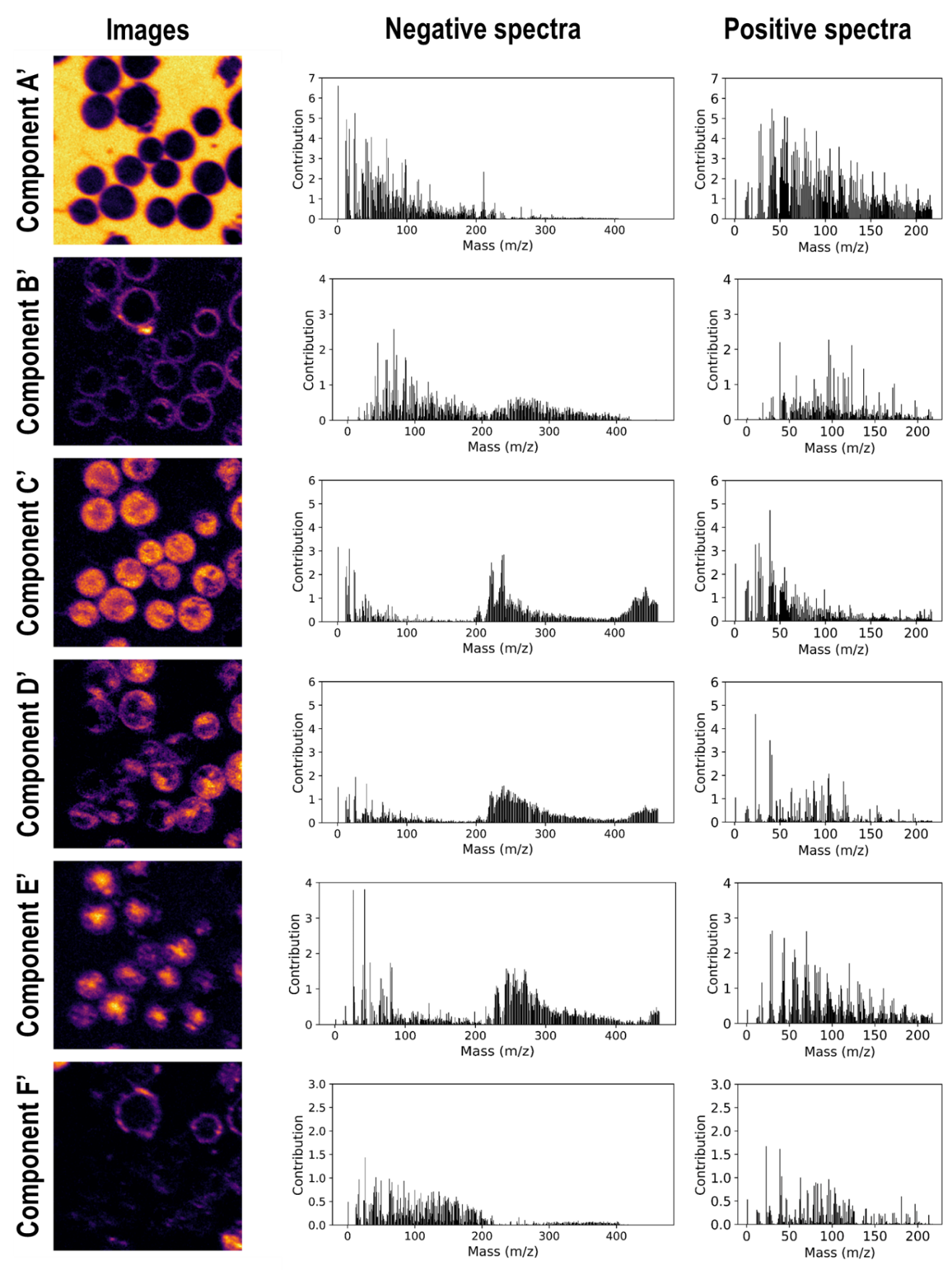

**Table S1**

Summary of analysis conditions for the 4 ToF-SIMS datasets presented in this study.

| Analysis | HLR - | HLR + | HMR - | HMR+ |
| --- | --- | --- | --- | --- |
| Pulse duration | 100 ns | | 15 ns | |
| Raster size | 100 x 100 µm | | | |
| Number of frames | 70 | 68 | 200 | 200 |
| Pixel number | 1024 x1024 | | 512 x 512 | |
| Pixel size (nm) | 97 nm | | 195 nm | |
| Lateral resolution measured on total ion image | 150 nm | 120 nm | 400 nm | 500 nm |
| Mass resolution | < 400 | < 400 | 9800 at mass 97  9260 at mass 189 | 9030 at mass 97  10390 at mass 211 |
| Peaks used for calibration | CN^-^, CNO^-^, PO_3_^-^ | CH^+^, Na^+^, C_7_H_7_^+^ | C_2_H^-^, C_3_H_3_-, C_4_H^-^, C_5_H_5_^-^, C_6_H_7_^-^, OsO_3_^-^ | C_2_H_2_^+^, C_3_H_4_^+^, C_5_H_5_^+^, C_7_H_6_^+^ |
| Mass deviation from calibration | -26.24 ppm | -4.42 ppm | 0.01 ppm | 0.04 ppm |
| Number of selected peaks | 509 | 255 | 761 | 488 |

**Table S2**

Identification of the top 20 contributors of component B using the correlative HLR/HMR approach.

| Order | Contribution | Polarity | Unit mass | Exact mass | Annotation | Mass dev (mamu) | Relative intensity (%) |
| --- | --- | --- | --- | --- | --- | --- | --- |
| 1 | 3.71 | - | 59 | 59.0136 | C_2_H_3_O_2_^-^ | 0.32 | 85 |
| 2 | 3.70 | - | 69 | 68.9972 | C_3_HO_2_^-^ | -0.47 | 62 |
| 3 | 3.64 | - | 45 | 44.9979 | CHO_2_^-^ | 0.06 | 79 |
| 4 | 3.63 | - | 41 | 41.0028 | C_2_OH^-^ | 0.04 | 88 |
| 5 | 3.37 | - | 87 | 87.0089 | C_3_H_3_O_3_^-^ | 0.75 | 42 |
| 6 | 3.33 | - | 71 | 71.0131 | C_3_H_3_O_2_^-^ | -0.13 | 86 |
| 7 | 3.12 | + | 99 | 98.9199 |  |  |  |
| 8 | 3.11 | - | 97 | 97.0284 | C_5_H_5_O_2_^-^ | -0.47 | 10 |
| 9 | 2.83 | - | 93 | 92.9268 |  |  |  |
| 10 | 2.75 | - | 99 | 99.0097 | C_4_H_3_O_3_^-^ | 1.35 | 66 |
| 11 | 2.50 | - | 75 |  |  |  |  |
| 12 | 2.49 | - | 83 | 83.0138 | C_4_H_3_O_2_^-^ | 0.59 | 41 |
| 13 | 2.48 | - | 67 | 68.9972 | C_3_HO_2_^-^ | -0.47 | 62 |
| 14 | 2.46 | - | 58 | 58.0053 | C_2_H_2_O_2_^-^ | -0.11 | 76 |
| 15 | 2.38 | - | 65 | 65.0019 | C_4_OH^-^ | -0.81 | 48 |
| 16 | 2.31 | - | 82 | 82.0047 | C_5_H_2_O_2_^-^ | -0.75 | 33 |
| 17 | 2.28 | - | 43 | 43.0188 | C_2_H_3_O^-^ | 0.39 | 88 |
| 18 | 2.27 | - | 86 |  |  |  |  |
| 19 | 2.23 | - | 73 | 72.9927 | C_2_HO_3_^-^ | 0.16 | 12 |
| 20 | 2.17 | - | 95 | 95.0130 | C_5_H_3_O_2_^-^ | -0.26 | 44 |

**Table S3**

Identification of the top 20 contributors of component C using the correlative HLR/HMR approach.

| Order | Contribution | Polarity | Unit mass | Exact mass | Annotation | Mass dev (mamu) | Relative intensity (%) |
| --- | --- | --- | --- | --- | --- | --- | --- |
| 1 | 4.04 | + | 41 | 41.0400 | C_3_H_5_^+^ | 0.94 | 73 |
| 2 | 3.18 | + | 43 | 43.0195 | C_2_H_3_O^+^ | 1.16 | 40 |
|  |  |  |  | 43.0562 | C_3_H_7_^+^ | 1.43 | 47 |
| 3 | 3.16 | - | 240 | 239.9477 | ^191^OsO_3_^-^ | 1.47 | 100 |
| 4 | 3.06 | - | 1 | 1.0081 | H^-^ | 0.28 | 100 |
| 5 | 3.05 | + | 55 | 55.0197 | C_3_H_3_O^+^ | 1.29 | 32 |
|  |  |  |  | 55.0562 | C_4_H_7_^+^ | 1.46 | 58 |
| 6 | 2.96 | + | 29 | 29.0034 | CHO^+^ | 0.65 | 15 |
|  |  |  |  | 29.0401 | C_2_H_5_^+^ | 0.97 | 76 |
| 7 | 2.65 | + | 57 |  |  |  |  |
| 8 | 2.62 | - | 238 | 237.9469 | ^190^OsO_3_^-^ | 3.70 | 100 |
| 9 | 2.55 | - | 222 | 221.9526 | ^190^OsO_2_^-^ | 4.30 | 90 |
| 10 | 2.53 | - | 237 | 236.9491 | ^189^OsO_3_^-^ | 6.20 | 88 |
| 11 | 2.46 | - | 13 | 13.0074 | CH^-^ | -0.43 | 100 |
| 12 | 2.45 | - | 224 | 223.9512 | OsO_2_^-^ | -0.07 | 84 |
| 13 | 2.31 | - | 25 | 25.0073 | C_2_H^-^ | -0.57 | 100 |
| 14 | 2.27 | + | 39 | 38.9646 | K^+^ | 0.87 | 81 |
| 15 | 2.26 | + | 1 | 1.0077 | H^+^ | -0.17 | 100 |
| 16 | 2.19 | - | 221 | 220.9539 | ^189^OsO_2_^-^ | -5.9 | 84 |
| 17 | 2.18 | + | 31 | 31.0192 | CH_3_O^+^ | 0.84 | 100 |
| 18 | 2.17 | - | 236 | 235.9459 | ^188^OsO_3_^-^ | -5.3 | 92 |
| 19 | 2.16 | - | 12 | 12.0003 | C | 0.29 | 100 |
| 20 | 2.15 | - | 17 | 17.0028 | OH^-^ | 0.04 | 100 |

**Table S4**

Identification of the top 20 contributors of component D using the correlative HLR/HMR approach.

| Order | Contribution | Polarity | Unit mass | Exact mass | Annotation | Mass dev (mamu) | Relative intensity (%) |
| --- | --- | --- | --- | --- | --- | --- | --- |
| 1 | 3.27 | - | 26 | 26.0036 | CN^-^ | 0.50 | 100 |
| 2 | 3.06 | - | 42 | 41.9985 | CNO^-^ | 0.51 | 92 |
| 3 | 2.62 | + | 70 | 70.0674 | C_4_H_8_N^+^ | 1.80 | 44 |
| 4 | 2.41 | + | 44 | 44.0508 | C_2_H_6_N^+^ | 0.84 | 73 |
| 5 | 2.18 | + | 30 | 30.0354 | CH_4_N^+^ | 1.09 | 56 |
| 6 | 1.96 | + | 68 | 68.0516 | C_4_H_6_N^+^ | 1.60 | 27 |
| 7 | 1.82 | + | 59 | 59.0719 | C_3_H_9_N^+^ | -1.69 | 38 |
| 8 | 1.75 | + | 86 | 86.0992 | C_5_H_12_N^+^ | 2.25 | 32 |
| 9 | 1.74 | + | 82 | 82.0684 | C_5_H_8_N^+^ | 2.74 | 46 |
| 10 | 1.73 | + | 80 | 80.0531 | C_5_H_6_N^+^ | 3.10 | 23 |
| 11 | 1.70 | - | 256 |  |  |  |  |
| 12 | 1.66 | + | 84 | 84.0833 | C_5_H_10_N^+^ | 2.05 | 55 |
| 13 | 1.60 | + | 42 | 42.0353 | C_2_H_4_N^+^ | 0.90 | 43 |
| 14 | 1.54 | - | 248 |  |  |  |  |
| 15 | 1.50 | - | 270 |  |  |  |  |
| 16 | 1.48 | + | 54 | 54.0354 | C_3_H_4_N^+^ | 1.01 | 17 |
| 17 | 1.47 | - | 66 | 65.9981 | C_3_NO^-^ | 0.14 | 54 |
| 18 | 1.46 | - | 258 |  |  |  |  |
| 19 | 1.43 | - | 269 |  |  |  |  |
| 20 | 1.41 | + | 96 |  |  |  |  |

**Table S5**

Identification of the top 20 contributors of component E using the correlative HLR/HMR approach.

| Order | Contribution | Polarity | Unit mass | Exact mass | Annotation | Mass dev (mamu) | Relative intensity (%) |
| --- | --- | --- | --- | --- | --- | --- | --- |
| 1 | 1.69 | + | 39 | 38.9646 | K^+^ | 0.87 | 81 |
| 2 | 1.39 | - | 253 |  |  |  |  |
| 3 | 1.29 | - | 243 | 242.9625 |  |  |  |
| 4 | 1.21 | - | 250 | 249.9545 |  |  |  |
| 5 | 1.15 | - | 241 |  |  |  |  |
| 6 | 1.13 | + | 23 | 22.9905 | Na^+^ | 0.79 | 100 |
| 7 | 1.12 | - | 247 |  |  |  |  |
| 8 | 1.10 | - | 262 | 261.9522 |  |  |  |
| 9 | 1.09 | - | 254 | 253.9603 |  |  |  |
| 10 | 1.09 | - | 255 | 254.9658 |  |  |  |
| 11 | 1.08 | - | 269 | 268.9625 |  |  |  |
| 12 | 1.06 | - | 248 | 247.9577 |  |  |  |
| 13 | 1.05 | - | 267 | 266.9663 |  |  |  |
| 14 | 1.05 | - | 265 |  |  |  |  |
| 15 | 1.04 | - | 245 |  |  |  |  |
| 16 | 1.04 | - | 261 | 260.9576 |  |  |  |
| 17 | 1.02 | - | 256 |  |  |  |  |
| 18 | 1.02 | - | 283 |  |  |  |  |
| 19 | 1.01 | - | 251 | 250.9649 |  |  |  |
| 20 | 1.01 | - | 278 |  |  |  |  |

**Table S6**

Identification of the top 20 contributors of component F using the correlative HLR/HMR approach.

| Order | Contribution | Polarity | Unit mass | Exact mass | Annotation | Mass dev (mamu) | Relative intensity (%) |
| --- | --- | --- | --- | --- | --- | --- | --- |
| 1 | 3.27 | + | 39 | 38.9646 | K^+^ | 0.91 | 81 |
| 2 | 3.23 | + | 23 | 22.9905 | Na^+^ | 0.76 | 100 |
| 3 | 1.85 | - | 17 | 17.0028 | OH^-^ | 0.04 | 100 |
| 4 | 1.80 | - | 1 | 1.0084 | H^-^ | 0.59 | 100 |
| 5 | 1.71 | - | 24 | 23.9997 | C_2_^-^ | -0.29 | 100 |
| 6 | 1.64 | - | 25 | 13.0080 | C_2_H^-^ | 0.20 | 100 |
| 7 | 1.63 | - | 13 | 13.0074 | CH^-^ | -0.43 | 100 |
| 8 | 1.57 | - | 12 | 12.0003 | C^-^ | 0.29 | 100 |
| 9 | 1.44 | - | 16 | 15.9949 | O^-^ | 0.03 | 100 |
| 10 | 1.30 | + | 41 | 40.9630 | ^41^K^+^ | 1.20 | 19 |
| 11 | 1.25 | + | 63 | 62.9831 | C_2_ONa^+^ | -1.56 | 26 |
| 12 | 1.05 | + | 1 | 1.0076 | H^+^ | -0.21 | 100 |
| 13 | 1.05 | - | 26 | 26.0036 | CN^-^ | 0.50 | 100 |
| 14 | 1.01 | + | 46 | 45.9804 |  |  |  |
| 15 | 0.98 | - | 48 | 47.9990 | C_4_^-^ | -1.00 | 93 |
| 16 | 0.96 | - | 40 |  |  |  |  |
| 17 | 0.95 | - | 36 | 35.9995 | C_3_^-^ | -0.55 | 100 |
| 18 | 0.92 | + | 88 | 87.9568 | CNNaK^+^ | 0.23 | 63 |
| 19 | 0.87 | + | 62 | 61.9546 | NaK^+^ | 1.13 | 23 |
| 20 | 0.87 | - | 14 | 14.01258 | CH_2_^-^ | 0.14 | 100 |
